# The Effect of Oral Tactile Sensitivity on Texture Perception and Mastication Behavior in Humans

**DOI:** 10.1101/466342

**Authors:** Grace E. Shupe, Arran Wilson, Curtis R. Luckett

**Affiliations:** Department of Food Science, Institute of Agriculture, University of Tennessee, Knoxville, Tennessee; The New Zealand Institute for Plant & Food Research Limited, Lincoln, New Zealand

**Keywords:** mastication, sensitivity, texture, discrimination

## Abstract

Mastication behavior is a notable source of interindividual variation in texture perception and could be linked to oral sensitivity. As oral sensitivity declines so does the amount of tactile feedback relayed to the brain, resulting in less effective manipulation or food and a reduced ability to discriminate differences. To address these hypotheses, we measured masticatory behavior and related this to texture discrimination and oral sensitivity. The study was performed on 41 participants in two groups, with high (n = 20) or low (n=21) sensitivity. Oral sensitivity was measured using a battery of tests that included: oral stereognosis, lingual tactile acuity, and bite force sensitivity. Sensitivity to texture changes was measured using a series of triangle tests with confectionaries of different hardness, with masticatory patterns and behaviors being video recorded and analyzed using jaw tracking software. Overall, there was no significant difference between high and low sensitivity participants and their ability to distinguish texture changes. But, there were significantly different trends found between the groups based on their masticatory behaviors including chewing pattern and overall number of chews. But, it was found that multiple masticatory behaviors were being modulated by oral sensitivity, including overall chewing cycles used (p < 0.0001). More, specifically those in the high sensitivity group used more stochastic chewing movements, while those in the low sensitivity group were found to use crescent-shaped chewing cycles. It was also noted that in the high sensitivity group the jaw moved further distances (p < 0.0001) in all phases and moved at a higher velocity when opening (p < 0.0001) but not when closing, when compared to the low sensitivity group. These results help bolster evidence that mastication and oral sensitivity are related.

## 1.1 Introduction

Texture perception is a dynamic process that is constantly changing during oral processing; therefore, mastication and texture perception are thought to be linked (Hutchings & Lillford, 1988). During mastication, the first step in the digestive system, a product is broken down in the oral cavity and its texture properties are continuously changing (e.g. particle size reduction, saliva lubricating and softening, mixing) (Hutchings & Lillford, 1988; Szczesniak, 2002). As feedback on these textural properties is received from the oral cavity adjustments are made to maximize the efficiency of chewing, altering masticatory patterns as well. This feedback from the oral cavity to the brain creates a loop that modulates force, energy, speed, etc. required to properly masticate and form a bolus (Lund, Kolta, Westberg, & Scot, 1998; van der Glas, van der Bilt, Abbink, Mason, & Cadden, 2007). The action of chewing is controlled by a central pattern generator located in the brainstem, modulating peak amplitudes, force loads and rhythmic movements (Avivi-Arber, Martin, Lee, & Sessle, 2011; Lund et al., 1998; Widmer & Morris-Wiman, 2018). Furthermore, people eat differently and have different mechanism for chewing, resulting in notable differences between consumers making it difficult to collect and compare behavior results. Because of this complexity, relatively few published papers have been published investigating the relationship of masticatory behavior and texture perception. In addition, many of the most comprehensive findings on the relationship between masticatory behavior and texture perception were primarily concerned with age related changes in either variable (Forde & Delahunty, 2002; Kremer, Mojet, & Kroeze, 2007). A preliminary study by Pedroni-Pereira et al., looking at objective and subjective (by means of a questionnaire) measures of masticatory function found no correlation between objective and subjective measures. The objective measures of masticatory function were maximum bite force and two measures of masticatory performance, all of these measures were moderately correlated (2018).

Oral sensitivity has been well documented to decrease with age, along with other forms of mastication performance such as chewing efficiency. Uniquely mastication, is key to many food sensations such as texture perception (Brown, Langley, Martin, & MacFie, 1994; Wilkinson, Dijksterhuis, & Minekus, 2000), flavor release (Taylor & Roozen, 1996), flavor perception (Luckett, Meullenet, & Seo, 2016), and bolus formation (Devezeaux de Lavergne, Derks, Ketel, de Wijk, & Stieger, 2015). All of this requires the active breakdown and manipulation of a food product in the oral cavity (Brown et al., 1994; Forde & Delahunty, 2002). It is thought that as oral sensitivity declines there will be less feedback from the oral cavity to the brain resulting in a less efficient and longer masticatory process. This could result in the use of compensatory strategies used by older populations, such as chewing longer or chewing more in a specified amount of time (K. Kohyama, Mioche, & Martin, 2002; Mioche, Boundial, & Peyron, 2004).

In general, people have different chewing styles and chewing efficiencies which can lead to different chewing times and swallowing thresholds(Brown et al., 1994; Devezeaux de Lavergne et al., 2015). The variance of mastication measurements obtained is relatively high due to intra-individual differences exhibited by participants, in a study by Remijn et al. chewing duration and chewing frequency showed the best reproducibility while chewing side and other measures were not reproducible when using 3D kinematics and sEMG (2016). It has been shown that chewing time can change a consumer’s perception of a food product since it is not manipulated for a long duration a soft product will be perceived as harder due to the breakdown process not being fully completed (Brown et al., 1994). Resulting in the first characteristics of a product being used for judgements by a fast eater, since later sensory information that a slow eater would have is unavailable to a fast eater (Brown et al., 1994). When comparing slow and fast eaters using soft and hard sausages, it was noted that there was difference in bolus properties at the end of mastication for these two groups (Devezeaux de Lavergne et al., 2015). However, using Temporal Dominance of Sensations (TDS), the first dominate attribute was not different between the fast and slow eaters. Conversely, the attributes did become different between the two groups towards the end of the mastication sequence.

Work has been going on for years on how to link subjective measures (such as those received from a sensory panel during Qualitative Descriptive Analysis (QDA) or TDS) to objective instrumental measurements of food texture properties (James, 2018; Le Révérend, Saucy, Moser, & Loret, 2016). In a study by Révérend et al., seven cereal products were used that had similar fracture force, all of which were perceived differently due to the internal structure (low density/high porosity) of each product (2016). Finding like these highlight why it is why it is so important to use a human observation to translation sensory perception to physical information such as that obtained from texture profile analysis (TPA)(James, 2018; Nishinari, Kohyama, Kumagai, Funami, & Bourne, 2013). Even using a model food stuff such a gel or agar, there will be melting and saliva incorporation during the end of oral processing and these factors cannot be recreated during TPA.

It would be logical for some of the individual variation in texture perception to be explained by differences in oral sensitivity. Kremer et al. (2007) reported a mild association between oral tactile sensitivity and texture perception, using chewing efficiency of two-color gum, oral stereognosis, and particle size discrimination; olfactory ability was also characterized for the elderly group only. Elderly participants were found to preform significantly worse at the chewing and oral stereognosis tasks but were not different in their ability to distinguish particle size when given two samples and asked to identify the finer sample. However, this study was not solely designed to characterize the relationship between oral sensitivity and texture perception, therefore several confounding factors make definitive conclusions difficult. For example, flavor preferences were based on participants olfactory acuity, the participants groups were split at the median, and the experimental groups had a relatively small *n*=10 and *12* for good and poor performers, respectively (Kremer et al., 2007). Forde and Delahunty (2002) showed that texture attributes were more important for liking in older participants than in younger participants when looking at liquid, semi-solid, and solid foods. Kremer et al. (2007) investigated the relationship of texture and flavor manipulation with sweet and savory waffles in young and old populations.

It was found that older populations had a decreased sensitivity to oral stereognosis but not when discriminating particles sizes; and older populations also exhibited lower chewing efficiency. This agrees with previous research, that not all sensations are influenced the same way during aging. Calhoun et al. (1992) found that vibration and thermal sensations were intact in older populations while two-point discrimination and oral stereognosis showed declines. Furthermore, when looking at the effects of mastication on food intake, where chewing cycles was modified to 100%, 150% and 200% of participants normal chews, younger participants had a 10% and 14% decrease in food intake but older participants had no such decline (Hollis, 2018; Zhu & Hollis, 2014).

The purpose of this study is to better understand the relationships between masticatory behavior and oral sensitivity. More specifically, to look for changes in masticatory behavior with difference in oral sensitivity. Secondarily, to quantify the relationship of texture perception and oral sensitivity. Hence, we hypothesize the following:

H_1_ High oral sensitivity participants will be more sensitive to texture difference between samples.

H_2_ Oral sensitivity will modulate masticatory behavior.

## 1.2 Materials and Methods

### 1.2.1 Participants

41 participants were recruited for this study. Participants reported no olfactory dysfunction, had no allergies or food restrictions, and were not pregnant. Participants were asked to self-report common dental procedures such as root canals, crowns, partial or full dentures. Participants were recruited by their oral sensitivity. Using the test battery outlined in Shupe et al. 2018, in which participants were characterized oral stereognosis, raised and recessed shape identification, and bite force sensitivity. The results of these three measures were compiled and a total score was calculated. This study recruited subjects that scored in the upper 25% of oral sensitivity and those that scored in the lower 25% of oral sensitivity. The high sensitivity group contained 20 participants, while the low sensitivity group was comprised of 21 participants (see Table 1 for participant demographics). All participants signed an informed consent and were compensated for their time. This experiment was conducted according to the Declaration of Helsinki for studies on human subjects and approved by the University of Tennessee IRB review for research involving human subjects (IRB #18-04466-XP). The authors declare that they do not have any conflict of interest.

**Table 1.**
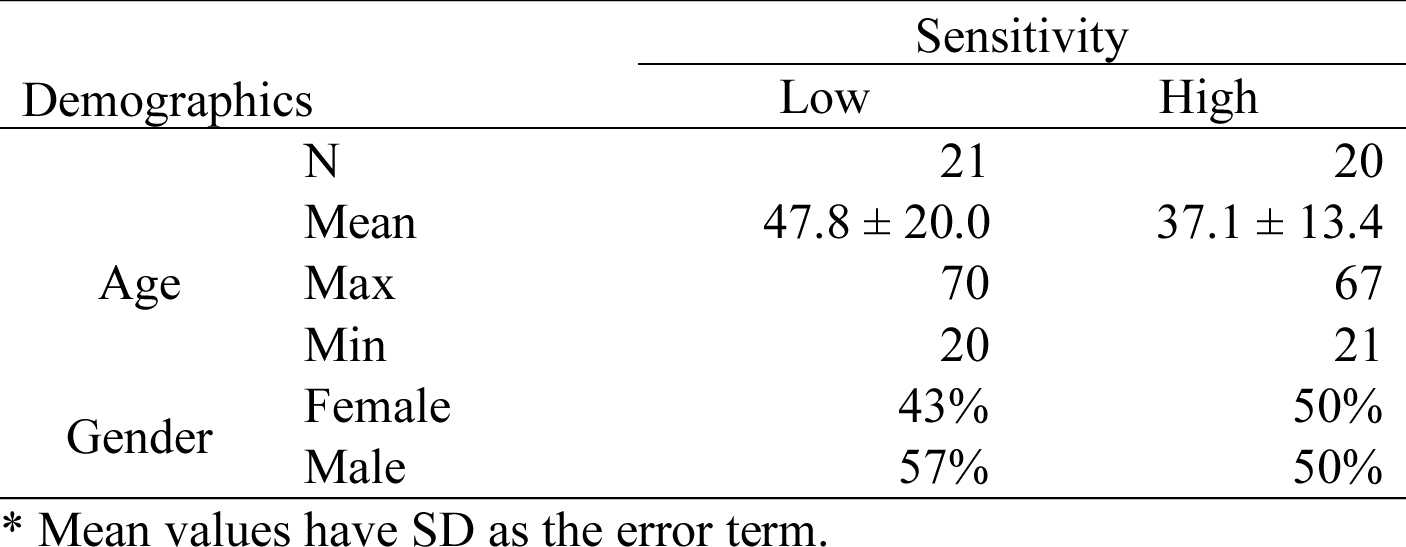
Demographics of participants by age group.

### 1.2.2 Stimuli

#### 1.2.2.1 Oral Sensitivity

Oral sensitivity stimuli were previously defined in Shupe et al., and consisted of oral stereognosis, lingual tactile sensitivity, and bite force sensitivity stimuli (2018).

#### 1.2.2.2 Confectionaries

Texture stimuli were made using the following ingredients, sucrose (Domino Foods Younkers, NY), glucose syrup (Caulet, Erquinghem-Lys, France), sorbitol (4mular, Irvine, CA), citric acid (SAFC, Switzerland), and water were mixed together and heated using a double boiling system until forming a homogenous solution (see Table 2). Three different gelatin bloom strengths were used to create texture differences (170, 200, and 230 bloom, Germany). Gelatin sheets were submerged in room temperature water until fully bloomed, approximately two minutes, then the gelatin was drained and added to the sugar solution; stirred using a stirring rod until completely dissolved, approximately two minutes. The solution was brought to room temperature (23 °C) and strawberry flavoring was incorporated. Then 4.0 g of the solution was poured into each oil coated, hemi-spherical silicone mold (11.2 cm^3^) and allowed to harden in a refrigerator (4°C) overnight. Confectionaries were verified at room temperature for hardness and springiness using a TA.XT Plus Texture Analyzer and Exponent software (Texture Technologies Corp. and Stable Micro Systems, Ltd., Hamilton, UK) shown in Table 3.

**Table 2.** Ingredients used to make confectionary texture stimuli.

| **Ingredient** | **Amount** |
| --- | --- |
| Sucrose | 200 g |
| Glucose Syrup | 300 g |
| Sorbitol | 15 g |
| Citric Acid | 6 g |
| Water | 232 g |
| Gelatin <sup>a</sup> | 7.5 g |
| Flavor <sup>a</sup> | 112 μl |

<sup>a</sup> Gelatin and flavor were both added to 1/8 (75 ml) of the sugar solution, in order to reduce variability of samples.

**Table 3.** Hardness of confectionary samples using TA.TX plus texture analyzer.

| Gummy | Bloom Strength | Hardness (N) ± SE | Springiness (%) ± SE |
| --- | --- | --- | --- |
| A | 170 | 2.54 ± 0.23 | 0.66 ± 0.04 |
| B | 190 | 2.10 ± 0.25 | 0.70 ± 0.03 |
| C | 200 | 1.68 ± 0.10 | 0.77 ± 0.03 |
| D | 230 | 1.75 ± 0.09 | 0.78 ± 0.02 |

#### 1.2.2.3 Jaw Tracking Apparatus

A polycarbonate face-shield (3M^™^, Saint Paul, MN, 55144) was transformed into an open front clear polycarbonate reference frame, similar to that used by Wilson et al. (2016). A quarter inch black reference line with a white boarder was visible from the front and a 1/8-inch diameter black dot surrounded by a white boarder was applied to each participant’s chin, this would allow software to track jaw movements (Figure 1).

**Figure 1.**
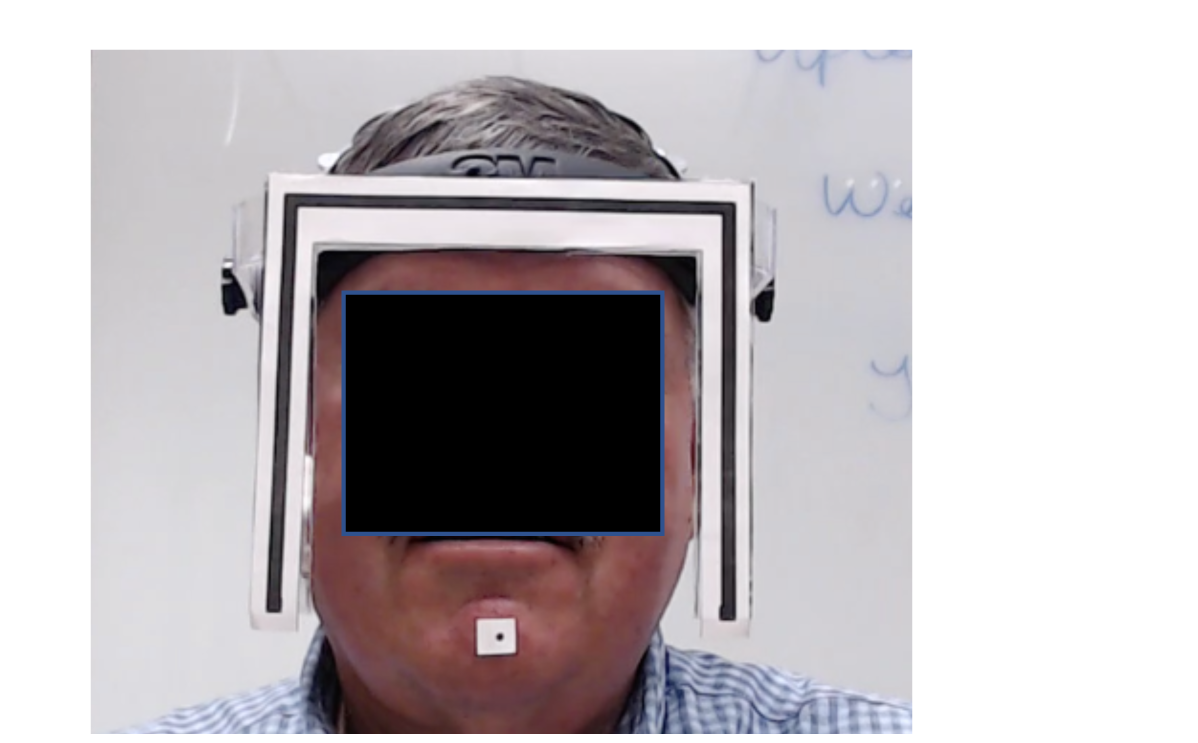
Example of head apparatus and chin marker that is used to track jaw movements.

### 1.2.3 Pre-Screening

The pre-screening process was completed as defined in Shupe et al. 2018, with participants receiving three different tests of oral sensitivity: oral stereognosis, raised and recessed shape identification, and bite force sensitivity. The results from Shupe et al. were used to determine high and low sensitivity, based on the distribution obtained from previous tests the upper and lower quartiles were used. Only those participants that were in the upper or lower quartile continued on to the jaw tracking exercise.

Dental status was self-reported by participants, categorized by 6 levels (*healthy or having filling(s)*, *singular crown*, *singular root canal procedure*, *multiple crowns and root canal procedures*, *dentures*, and minimal natural teeth with no prosthetics). Those subjects who reported *dentures* or *minimal natural teeth with no prosthetics* were considered notably compromised and were excluded from the study as dental status was not a factor of interest.

### 1.2.4 Procedure

During the first session, each participant was familiarized with each stimulus and the general tasks to be completed. The presentation of stimuli within each test was randomized, with the overall order of presentation maintained between participants to reduce fatigue. Participants completed one (1) approximately hour-long session with the following serving order; gummy letters (3), shapes, gummy letters (3), shapes, foam, gummy letters (3). Participants were asked to verbally respond with all answers, which were then recorded by members of the research team.

During the second and third sessions, participants were familiarized with video equipment used by the jaw tracking software and the discrimination task that they would be completing. Participants were also familiarized with the head apparatus that would be worn during testing and were instructed to place the entire sample in the mouth before chewing. They were also informed of the location of the camera and to look directly into it while chewing samples. Participants were then outfitted with the head apparatus and the chin dot before testing began. Participants were given three discrimination tasks, in order to determine sensitivity to texture changes. A triangle test was chosen as the discrimination task. In the triangle test, participants were presented three (3) samples simultaneously. Of these three, two are alike and one is different or “odd”. The participants are instructed to select the odd sample. This discrimination task was done in an unspecified manner, meaning the participants were not informed as to which characteristic (i.e. texture) differed amongst the samples. Through-out testing, participants filled out demographics; upon completion of the three sessions they were compensated for time participating.

### 1.2.6 Data Analysis

Data was structured as the number of correct responses each panelist gave for all oral sensitivity measures. In order to determine overall oral sensitivity, the sum of correct responses for lingual sensitivity, stereognosis, and bite force sensitivity tasks was used (Shupe et al. 2018).

Jaw tracking videos were recorded using a Logitech HD Pro Webcam C920. Each video consisted of the chewing sequence of a single sample from the discrimination task, therefore each participant produced nine (9) mastication sequences for analysis. The videos were then analyzed using the method defined by Wilson et. al (2016). This method uses the corner points of the tracking frame to define a facial plane, which is then transformed so that plane lies perpendicular to the viewer’s line of sight and is scaled to the dimensions of the frame. The location of the chin dot is then tracked within each transformed image giving a 2D chewing trajectory that is corrected for head movement thus allowing measurements such as vertical and horizontal distances, speeds, velocities, angles, and slopes etc to be made (see Table 4 for a list of all variables). Because the web cam frame rate varied around the nominal 25 frames/sec between recordings the actual frame rate was used to calculate all secondary measures, each individual frame rate from each video was used to calculate distance, speeds, and velocities of jaw movements these could be specified as jaw opening and closing. Also, chewing cycle shapes (circular, crossed, crescent, and no shape) were also determined as shown in Figure 2. These results of the video analysis were used to determine difference between groups. Two participants were excluded for masticatory behavior analyses due to the poor video quality, these two participants were included in discrimination analyses.

**Figure 2.**
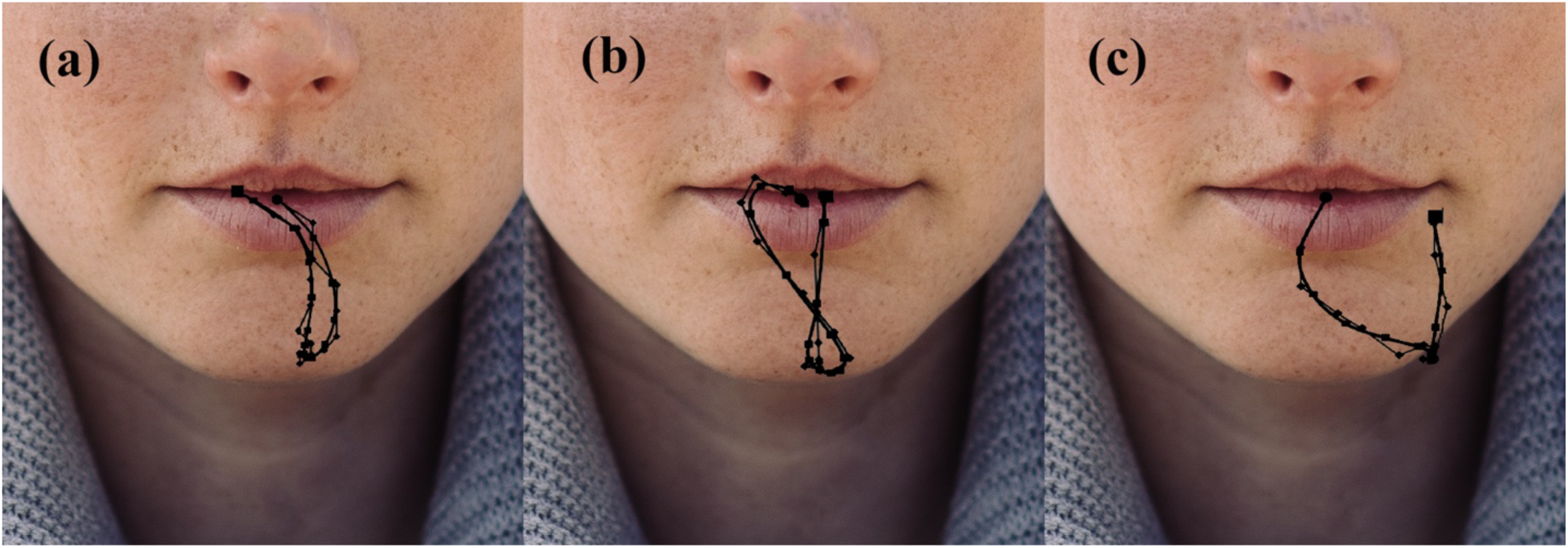
Examples of different known chewing patterns (a) crescent (b) crossed and (c) circular.

**Table 4.** List of variables extracted from the video clips.

| Variable Name | Description | Units |
| --- | --- | --- |
| Chews | Total number of chews | chews |
| Chew Time | Total chewing time | Seconds |
| Frequency | mean inverse of seconds per chew | chews/second |
| Close | mean closing distance | mm |
| CloseV | mean closing speed | mm/second |
| Open | mean opening distance | mm |
| OpenV | mean opening speed | mm/second |
| Width | mean width of chew | mm |
| Height | mean height of chew | mm |
| Perimeter Length | mean distance of chew | mm |
| Crossed Cycle | proportion of crossed shaped chews | - |
| Crescent Cycle | proportion of crescent shaped chews | - |
| Circle Cycle | proportion of circular shaped chews | - |
| No Shape Cycle | proportion of chews with no shape | - |

Results were analyzed using R and JMP Pro 13.1 (SAS Institute, Cary, NC), with statistically significant defined as p < 0.05. Differences in jaw movement and chewing parameters were examined across sensitivity level and sample by multiple analysis of variance (ANOVAs). In order to verify the assumptions of a T-test for high and low sensitivity groupings, variance was compared using Brown-Forsythe test of unequal variance. Six of the 20 variable had unequal variance between high and low sensitivity groupings, and a Welch’s t-test was preformed to account for the assumptions not being met. Pairwise post-hoc comparisons were performed using Tukey’s HSD adjustment and simple correlations were used to determine associations between measures. To compare masticatory behavior, multiple ANOVAs were run using sample and sensitivity group (high or low) as fixed factors. A multiple logistic regression model was run to identify which mastication behaviors lead to texture discrimination.

## 1.3 Results and Discussion

### 1.3.1 Age and Gender

There was not a significant difference in age between sensitivity groupings (*X*^2^_1_= 3.31, p = 0.07). There was also not a significant difference in sensitivity groupings between gender (*X*^2^_1_= 0.21, p = 0.65).

### 1.3.2 Sensitivity to Texture Changes

There was not a significant difference in sensitivity to texture changes between high and low sensitivity groupings (p = 0.486), showing that oral sensitivity does not have an effect on the texture discrimination of gummy confections (as shown in Figure 3). Overall participants performed well on this discrimination task, with approximately 40% of all participants correctly identifying the odd sample regardless of sensitivity grouping.

**Figure 3.**
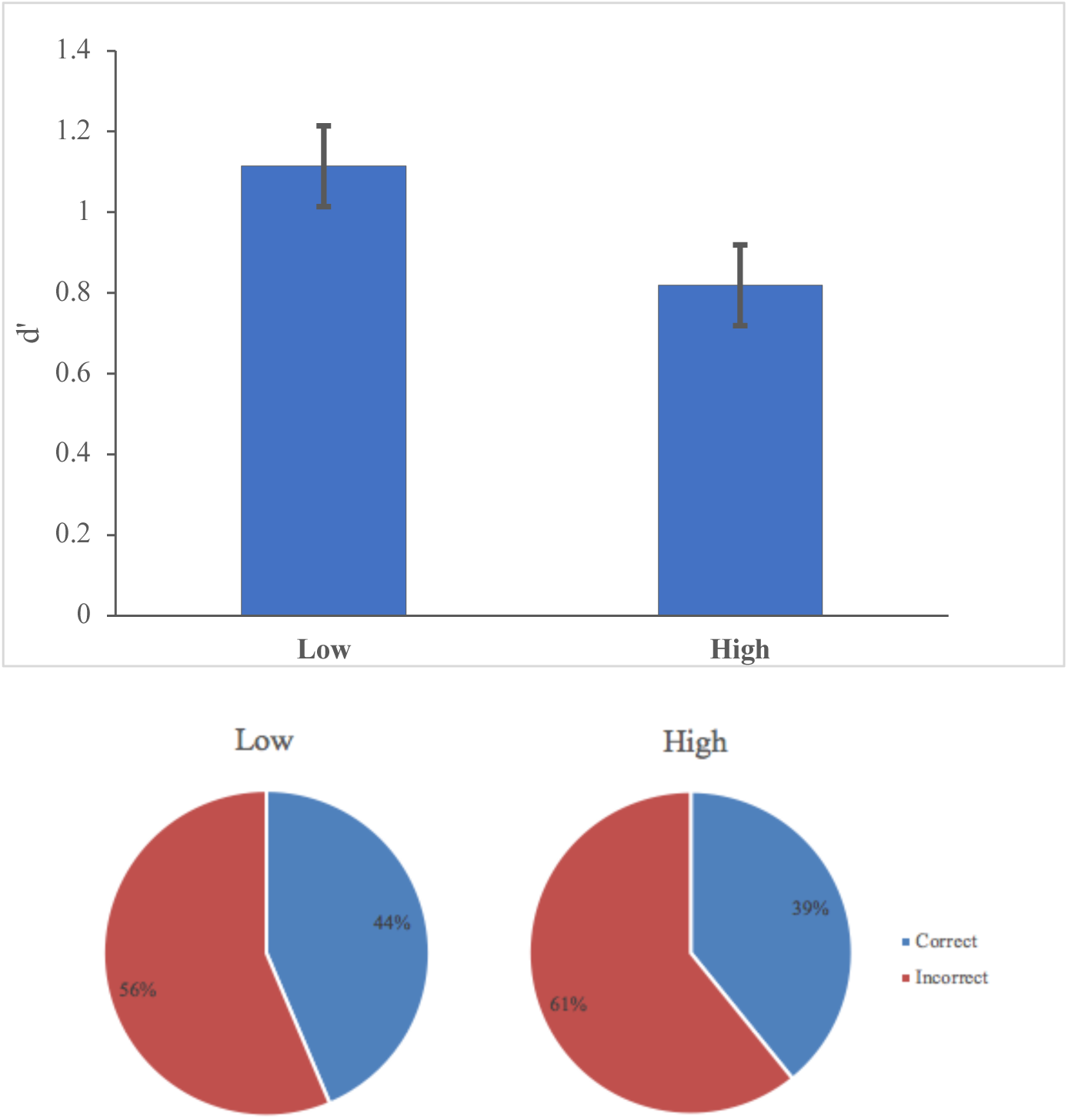
d’ for discrimination task for low and high sensitivity groups and proportions of correct responses.

The present results show that as oral sensitivity increases there is no corresponding increase in discrimination ability. It was expected that oral sensitivity would modulate a participant’s sensitivity to texture changes. As oral sensitivity increases, the potential for textural information from the oral cavity to the brain also increases, which can be clearly seen when prosthetics are used, since the removal of natural teeth drastically limits the information available to the masticatory feedback loop by excluding the tactile feedback from the periodontal ligament (which is the main provider of mechanical feedback in the oral cavity) (Trulsson, 2005). It has also been shown that even when physiological declines are present, there is not always a corresponding decline in sensory perception, especially when dealing with dynamic systems.

In the work by Kremer et al. (2007), it was noted that even when sensory declines of oral sensitivity are present in older populations there was not a related perception decrease of texture attributes. Furthermore, this was not the case when there was an olfactory sensitivity decline, this resulted in a decrease in flavor intensity ratings for sweet and savory waffles. It has also been documented that chewing behaviors can influence flavor and texture perception (Brown et al., 1994). Specifically those foods that are firm or rubbery as these were rated significantly different by groups that exhibited fast and slow eating behaviors (Brown et al., 1994).

The masticatory process is largely automatic when the task is easy, similarly to when you are walking through a familiar area (no extra though is needed to know where you need to go) (Lund et al., 1998; Ottenhoff, van der Bilt, van der Glas, & Bosman, 1992; van der Glas et al., 2007). But, when you are asked questions about a food product, there is more thought that goes into analyzing the components than normal masticatory patterns, providing enough difficulty in the discrimination task. However, the difference between each sample was too obvious to obtain clear separation based on oral sensitivity levels of each group. A harder discrimination task would give a wider range of ability between participants. Although no relationships were found between oral sensitivity level and sensitivity to texture changes it was noted that oral sensitivity is a significant factor in explaining the variance between masticatory behavior and chewing patterns.

### 1.3.3 Mastication Behavior

We were able to verify that the texture modifications received different oral processing and were different enough to extract different parameters even despite the ease of the discrimination task. As shown in Figure 4, as the hardness of the confectionary increase so does the total number of chews prior to swallowing (F_4, 654_ = 2.99, p = 0.0304).

**Figure 4.**
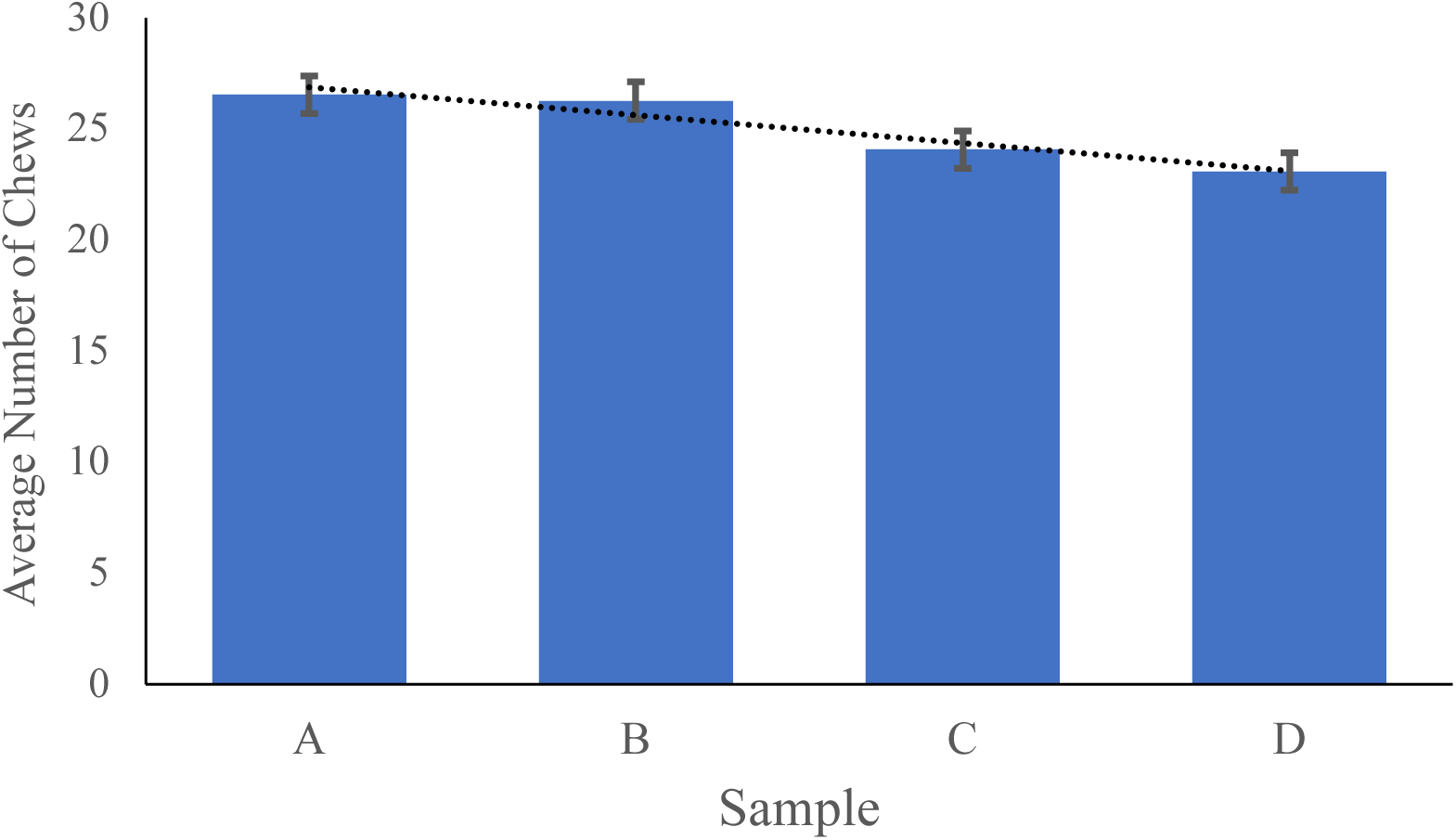
Total chews prior to swallowing for each confectionary hardness.

Overall masticatory behavior was different across the sensitivity groups with many of the variables having significant differences between the groups. In looking at specific chewing patterns used, crescent and crossed chewing patterns were significantly used more by the lower sensitivity group (F_1,656_= 11.86, p = 0.0006 and F_1,660_= 9.54, p = 0.0021 respectively), as shown in Figure 5. The high sensitivity group was significantly high in the amount of no shape chewing patterns used (F_1,656_= 22.16, p < 0.0001). Conversely, circular chewing patterns did not differ by age group (F_1,656_ = 0.04, p = 0.84). This shows the high sensitivity participants, when compared to low sensitivity participants, are much more likely to use novel or unpredictable chewing patterns based on the feedback that is received during chewing.

**Figure 5.**
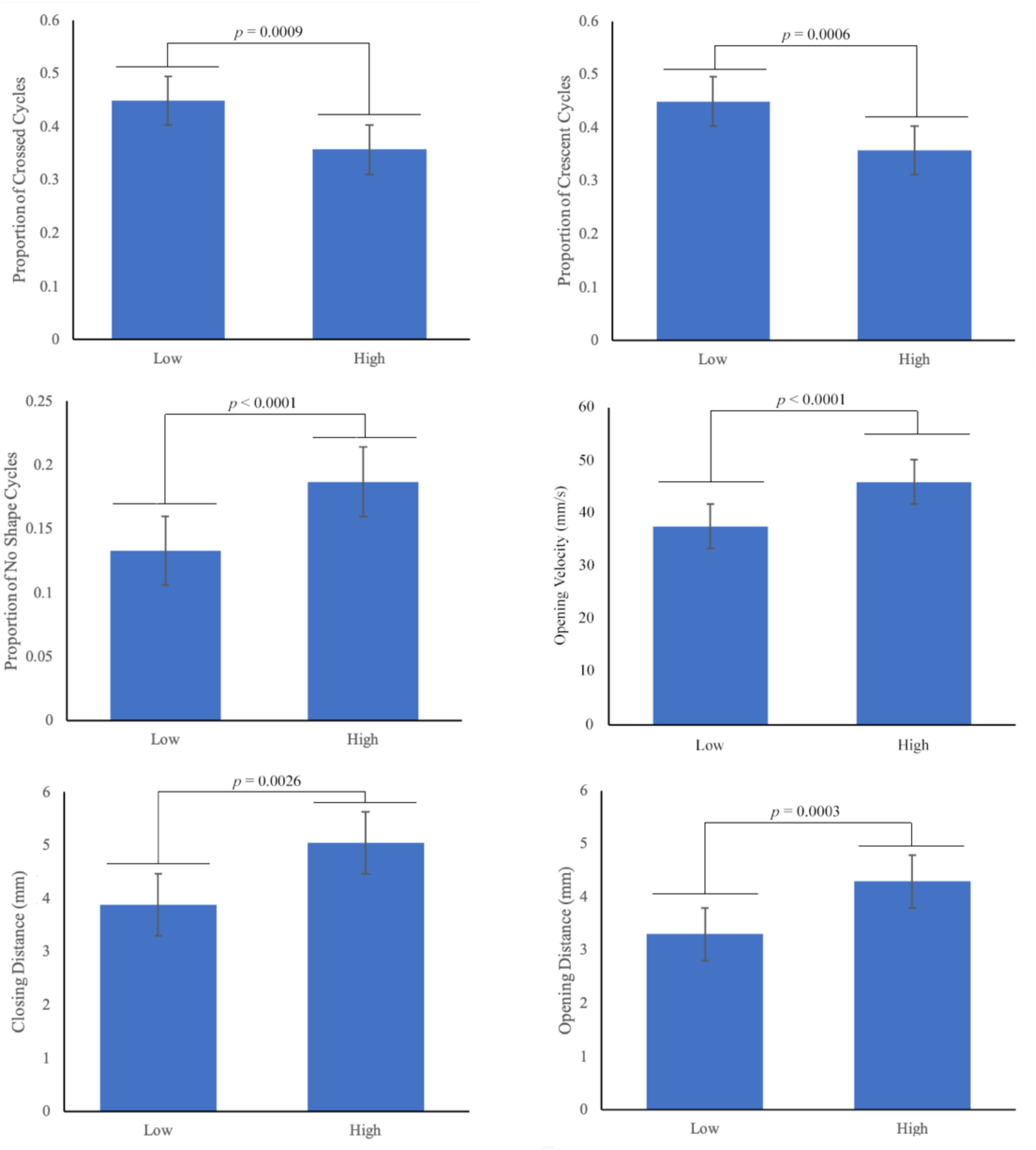
Mean values when comparing low and high oral sensitivity groups.

Further investigations into chewing parameters showed that there were significant differences between sensitivity group’s physiological measure of chewing, such as opening and closing distances. High sensitivity participants had a significantly larger opening and closing distance (F_1,656_ = 12.96, p = 0.0003 and F_1,656_ = 9.17, p = 0.0026, respectively), as shown in Figure 6. This results in a significantly larger average height and width of chew distance than the lower sensitivity group (F_1,656_ = 11.72, p = 0.0007, and F_1,656_ = 8.09, p = 0.0046, respectively). The overall perimeter chewing length was also significantly higher for the high sensitivity participants (F_1,656_ = 15.53, p < 0.0001), which is to be expected from the above findings. Therefore, considering all the above it would be expected that high sensitivity participants would be slower chewer, when in fact, high sensitivity participants were not significantly different from the lower sensitivity participants in both total number of chews and total chew time (F_1,656_= 0.05, p = 0.82 and F_1,656_ = 0.27, p = 0.61, respectively).

Overall, the frequency at which high and low sensitivity participants chew (chews/second) was not significantly higher for high sensitivity participants (F_1,556_= 0.68, p = 0.41). The mean opening and closing velocities noted for high sensitivity participants is significantly faster for opening, but not for closing, than that of the low sensitivity participants (F_1,657_ = 28.2, p < 0.0001 and F_1,657_ = 0.74, p = 0.39, respectively). On average a high sensitivity participant’s chewing would be described as more exuberant than low sensitivity participants, who have a slower more paced rhythmic chewing cycle, which can be confirmed by the higher proportion of known chewing patterns being used. The finding that high sensitivity participants are more active chewers agrees with previous research (Engelen, Van der Bilt, & Bosman, 2004; K. Kohyama, Mioche, & Bourdiol, 2003; K. Kohyama et al., 2002). It has been noted in older populations there is a decrease in oral sensitivity, which can lead to compensatory strategies such as chewing for longer periods of time or chewing more in a specified amount of time (K. Kohyama et al., 2002; Mioche et al., 2004). These same strategies seem to play a role whenever oral sensitivity is not optimal, regardless of age.

It is also of interest that low oral sensitivity participants have a higher variance, when compared to the high sensitivity participants. Three of the variables showed unequal variance by the Brown-Forsythe test. Both the mean opening slope (in degrees) showed significantly higher variance in the low oral sensitivity group when compared to the high sensitivity group (F_1,648_ = 7.41, p = 0.0067). It was also noted that chewing frequency variance was significantly different (F_1,657_ = 46.8, p < 0.0001). In both of these comparisons the low sensitivity group showed a lack of control for these parameters, resulting in large variations in values from this group. As previously mentioned, this lack of control could be a result of a lack of feedback from the oral cavity due to a loss in sensitivity, which would result in a lack of confidence and potentially slower jaw movements.

Confidence is commonly studied about sport and motor movements, but the same theory can be applied here. There are two factors involved when developing confidence in movements, this is competency of the jaw muscles themselves and movement sense (or the expected sensory experience when that muscle is moved) (Griffin & Keogh, 1982). Jaw muscles and teeth have to potential to do harm; therefore, these movements need to be closely monitored in order to be confident when chewing. If either a person’s ability or senses are lacking, this would result in a lack of movement confidence in the jaw bite (Griffin & Keogh, 1982). Since slower jaw closing velocities were noted in both groups, this is when potential damage could occur in the oral cavity (i.e. clashing of teeth, biting lips/tongue). When looking at the low sensitivity group they are significantly slower when opening the jaw as well, which has significantly few hazards when compared to closing showing a potential lack of confidence. The same speed increase can also be noted in cognitive experiments, in a study looking at decision speeds and reported confidence (Geller & Pitz, 1968). As participants became more confident in their decisions, their decision speed also increased.

It was also noted that masticatory behaviors were slight difference between sexes. Females overall, regardless of sensitivity level, had a faster opening velocity (F_1,660_ = 10.18, p = 0.0015), while males had a faster closing velocity when compared to females (F_1,660_ = 6.69, p = 0.0099). There was also an interesting trend with the low sensitivity females using significantly lower proportion of no shape chewing cycles (F_1,660_ = 24.43, p < 0.0001) and also a higher proportion of crossed chewing cycles when compared to all other groups (F_1,660_ = 50.64, p < 0.0001). Although these results are not conclusive due to the small sample size that each group (low sensitivity: male n =10 and female n=10; high sensitivity: male n=12 and female n = 9), these results are supported by previous work (de Oliveira Scudine et al., 2016). A study by de Oliveira Scudine et al. showed that males had a higher maximum bite force and depended more on their larger muscle capacity resulting in a higher masticatory performance; while for females masticatory performance was based on chewing cycle patterns and overall chewing frequency (2016). Another study by Kohyama et al., was conducted solely with males in order to account for these potential sex differences in masticatory behavior (2016).

### 1.3.4 Limitations

Participants were instructed on how to perform the task prior to recording, some previously discouraged actions were still preformed and had to be corrected through-out testing. Due to the nature of video recording, in order to keep mastication as normal as possible the researchers did not intervene during a discrimination task. This would result in a loss of more data through talking or other unnatural movements. Therefore, any modification that needed to be made to a participant’s behavior (i.e. moving hands from view, not swallowing between samples, etc.) were discussed between triangle testing. This resulted in some jaw tracking data not being able to be analyzed (e.g. chews were cut off or missed, the reference corners were not visible).

## 1.4 Conclusion

Our results show that there are notable differences between the masticatory behaviors of high and low oral sensitivity groups, but there no such relationship between sensitivity to texture changes and sensitivity level. The lower sensitivity group tended to have higher levels of intragroup variance in mastication parameters than the high sensitivity group. High sensitivity participants were also more likely to use novel chewing patterns based on the feedback that is obtained during oral processing. Further research is required to quantify the relationship between oral sensitivity and texture discrimination utilizing a more difficult discrimination task.

## 1.5 Acknowledgements

The authors would like to thank Anita Best for her assistance in measuring oral sensitivity.

## 1.6 Author Contributions

Author Shupe prepared experiments, collected and analyzed the data, and drafted the manuscript. Author Wilson analyzed and interpreted the mastication data. Author Luckett designed the study, interpreted the data, and edited the manuscript.

